## Supplementary_Materials_Figures for "Domain loss enabled evolution of novel functions in a gene superfamily"

1. **Dataset in a spreadsheet format, including aligned sequences, grouping and localization predictions (separate file)**
2. **TOP1MT region gene map figure**
3. **Phylogenetic trees**
    a) DALI
    b) iqTree
    c) Exabayes
4. **CLANS analysis**
5. **DBSCAN clusters**
6. **Structures in .pdb format (separate file)**
7. **TOLIP protein space clustering (separate file)**
8. **Protein language models' embeddings (separate file)**

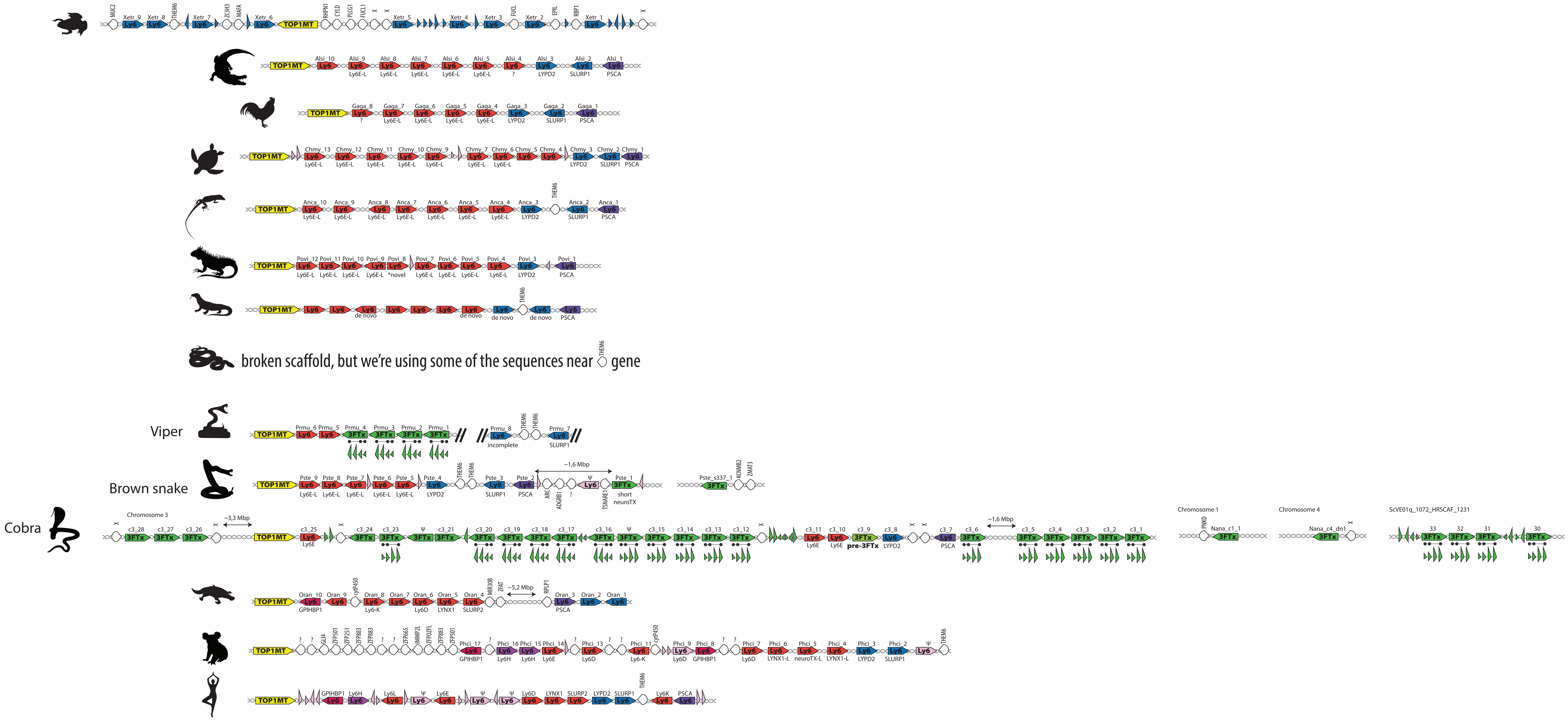


**Supplementary Material 2: Gene maps for TOP1MT genomic region of representative species.** Yellow arrow marks TOP1MT gene, other colored arrows mark Ly6 genes colored by orthology, white pointed semicircles mark any other gene present in the cluser (names given above). Triangles in line with the genes repre- sent orphan Ly6 exons colored by inferred homology. Triangles under 3FTx genes mark exonic structure of those genes that have non-canonical 4 exon structure, with a 3rd exon always pseudogenised. Names under the genes are given exactly as they are in genomic annotations. Names on top of the genes stand for the code used in our analysis.

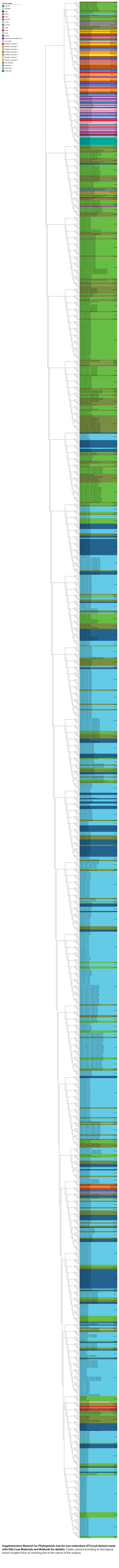

**Supplementary Material 3a: Phylogenetic tree for non-redundant 3FTx/Ly6 dataset made with DALI (see Materials and Methods for details).** Clades colored according to the legend, branch lengths have no meaning due to the nature of the analysis.

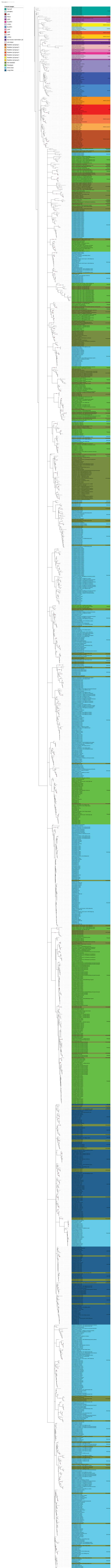


**Supplementary Material 3b: Phylogenetic tree for non-redundant 3FTx/Ly6 dataset made with iqTRee(see Materials and Methods for details).** Clades colored according to the legend, nodes with boostrap value higher than 85 are marked with a red dot.

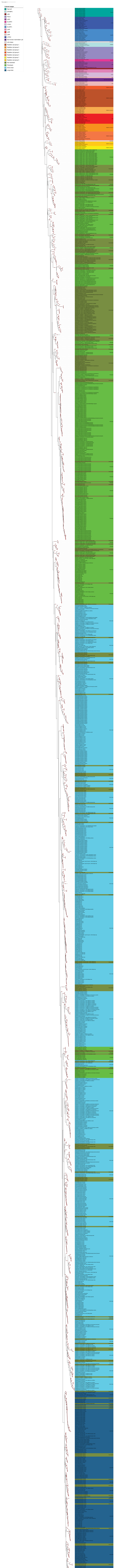


**Supplementary Material 3c: Phylogenetic tree for non-redundant 3FTx/Ly6 dataset made with ExaBayes (see Materials and Methods for details).** Clades colored according to the legend, nodes with posterior probability value higher than 85 are marked with a red dot.

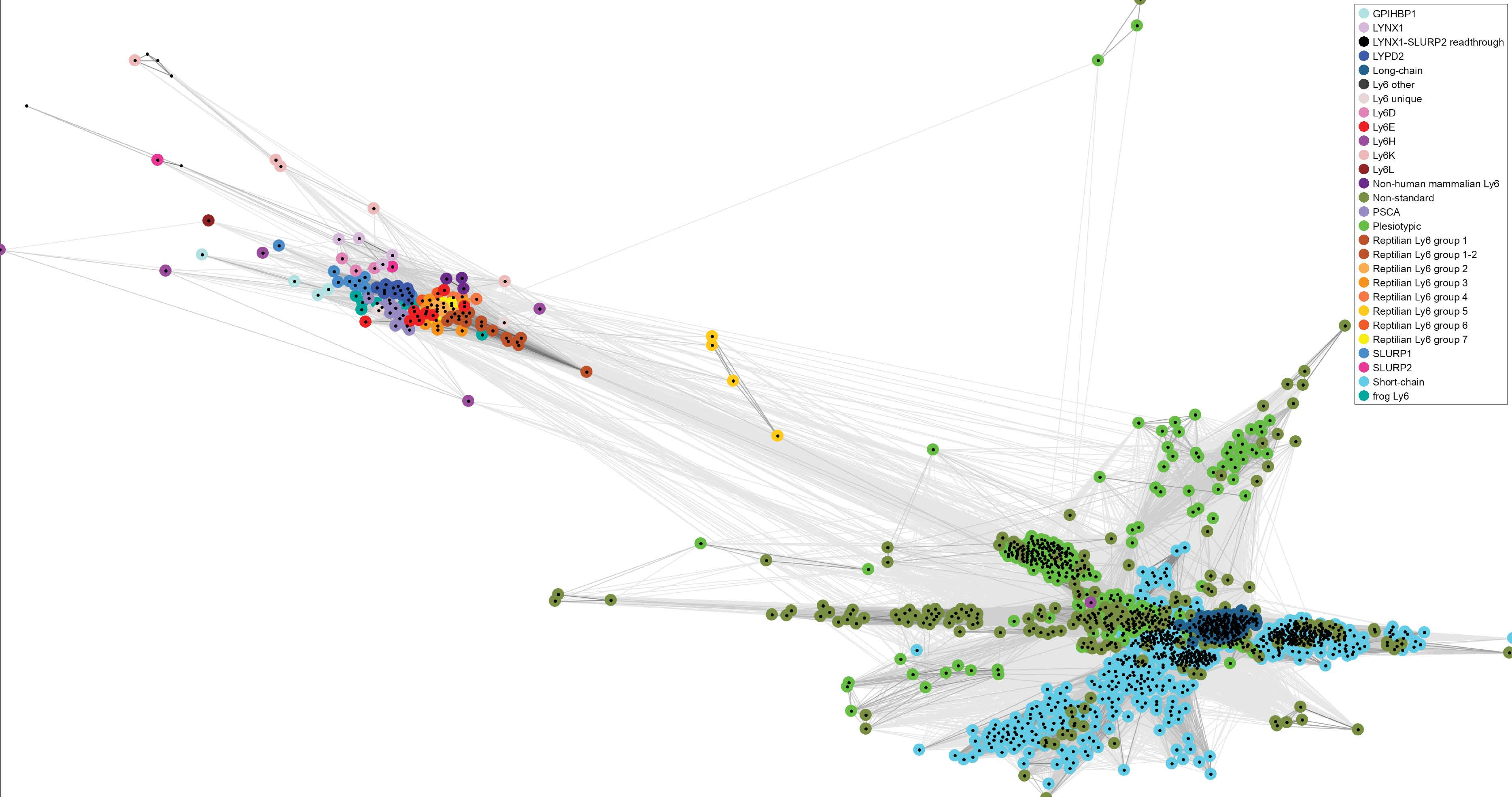


**Supplementary Material 4: CLANS sequence similarity based clustering for the non-redundant 3FTx/Ly6 dataset.** Datapoints colored according to the legend. The process involved an all-against-all BLAST comparison via the MPI Bioinformatics Toolkit's online CLANS tool. The image was produced using the CLANS Java application in 2D mode with 10,000 iterations, employing a p-value cutoff of > 1e-5.

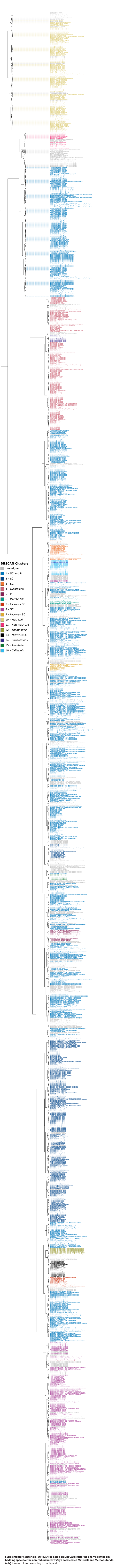


**Supplementary Material 5: OPTICS tree based on DBSCAN clustering analysis of the embedding space for the non-redundant 3FTx/Ly6 dataset (see Materials and Methods for details).** Leaves colored according to the legend.
